# *In utero* estrogenic endocrine disruption alters the stroma to increase extracellular matrix density and mammary gland stiffness

**DOI:** 10.1101/749044

**Authors:** Clarissa Wormsbaecher, Andrea R. Hindman, Alex Avendano, Marcos Cortes, Andrew Bushman, Lotanna Onua, Claire E. Kovalchin, Alina R. Murphy, Hannah L. Helber, Ali Shapiro, Kyle Voytovitch, Xingyan Kuang, Jonathan W. Song, Craig J. Burd

## Abstract

*In utero* endocrine disruption is linked to increased risk of breast cancer later in life. Despite numerous studies establishing this linkage, the long-term molecular changes that predispose mammary cells to carcinogenic transformation are unknown. Several lines of evidence indicate the stroma mediates endocrine disruption following early-life (or *in utero)* exposure. Herein, we utilized BPA as a model of estrogenic endocrine disruption to analyze the long-term consequences in the stroma. Using RNA-seq transcriptional profiling of adult primary fibroblasts isolated from female mice exposed to BPA *in utero*, we identified deregulated genes associated with the extracellular matrix. Specifically, multiple collagen genes had increased expression in exposed mice. In line with the transcriptional data, collagen deposition is increased in adult BPA-exposed mice. We further demonstrate *in vitro* that fibroblasts exposed to BPA *in utero* remodel a collagen matrix, thereby decreasing permeability of the collagen matrix. These alterations to the mammary gland resulted in increased gland stiffness in the adult mice. Our data connects early life endocrine disruption to breast density. Interestingly, increased collagen deposition and gland stiffness were not observed in the developing glands of younger mice, suggesting risk factors for breast cancer continue to develop throughout life following these exposures. Finally, we assessed whether *in utero* exposure to two other endocrine disruptors, BPS and DES, also increase breast stiffness in adult mice. While DES increased breast stiffness, BPS did not, suggesting this BPA alternative may in fact pose less breast cancer risk than its predecessor. As breast stiffness, extracellular matrix density, and collagen deposition have been directly linked to breast cancer risk, these data mechanistically link endocrine disruptor exposures and molecular alterations to increased disease susceptibility in the gland.

## Introduction

*In utero* exposure to estrogenic endocrine disruptors (EDCs) is associated with increased risk of breast cancer. Women exposed in the womb to diethylstilbestrol (DES) have twice the risk of developing the disease after the age of 40 ((NIH) 1999; Palmer et al. 2006). While *in utero* exposures to other endocrine disruptors is hard to quantify in the human population, animal models have demonstrated that many EDCs can also increase the risk of tumorigenesis. One of the most prevalent industrial EDCs is bisphenol A (BPA), which is used in plastics, canned food linings, thermal paper receipts, and other consumer products. Its abundant use has led to significant human exposures and is detected in human serum, urine, amniotic fluid, and fetal serum (Ikezuki et al. 2002; Schonfelder et al. 2002). The ability of BPA to act as an estrogen has raised concerns over these exposures, leading to its replacement in various commercial products. These BPA replacements have varying levels of estrogenic activity with compounds such as bisphenol AF having higher estrogenic activity than BPA and bisphenol S (BPS) having lower estrogenic activity (Kojima et al. 2019; Ng et al. 2015; Rochester and Bolden 2015). As the number of EDCs expands, understanding the critical properties and mechanisms driving risk becomes more imperative.

Rodent models demonstrate that early life exposures to estrogenic EDCs have long term consequences on mammary gland development and promote tumorigenesis. *In utero* BPA exposures induce preneoplastic lesions and hyperplasia in rat mammary glands (Durando et al. 2007; Murray et al. 2007). Mice exposed to BPA *in utero* develop tumors more quickly and more often when challenged with a chemical carcinogen (Weber Lozada and Keri 2011). Taken together, these rodent models show that early life BPA exposure increases the risk of mammary gland carcinogenesis. While BPA has been extensively studied for both developmental defects and cancer-associated phenotypes in rodent models, there is much less data on the BPA alternatives. However, BPS has been shown to drive many of the same developmental abnormalities in the mammary gland seen in BPA exposed animals (Kolla et al. 2018). While no study has demonstrated cancer formation in BPS-exposed mice, a small number of mice did have mammary neoplasias in one study (Tucker et al. 2018). These data suggest that compounds such as BPS are also a potential risk, yet no defined mechanism by which these compounds initiate oncogenesis has been identified. It is unclear whether the estrogenic properties of these EDCs are responsible for the increased cancer risk and which molecular alterations within the gland predispose the mammary cells to carcinogenic transformation. Without this knowledge, it is difficult to test and identify potentially hazardous compounds and account for latent effects following early life exposures.

As many of the EDCs that alter mammary gland development are estrogenic, the ability of these chemicals to stimulate the estrogen receptor have been used as the fundamental characteristic to identify them as potentially harmful compounds. Attempts to link the estrogenic action of BPA to the phenotypic changes in the mammary gland have implicated the developing mesenchyme as a potential cellular target of EDC action (Speroni et al. 2017). The developing mammary bud is completely surrounded by estrogen receptor (ER) expressing mesenchymal cells from embryonic day 12.5 to 15.5 (Hindman et al. 2017). This period during development appears to be the time point most susceptible to BPA action (Hindman et al. 2017). Furthermore, transcriptome profiling of the perinatal stroma and epithelium of the developing mammary gland at the time of exposure suggests that BPA is acting as an estrogen through the mesenchymal tissue (Paulose et al. 2015). This transcriptional profiling suggests that the extracellular matrix (ECM) is differentially regulated by BPA exposures (Paulose et al. 2015; Wadia et al. 2013). As growth, survival, and differentiation of the epithelium are dependent upon signals from the stroma (Macias and Hinck 2012), these alterations can influence carincogenic transformation later in life. To this end, we aimed to analyze the long-term changes within the fibroblast population that persist into adulthood when carcinogenic transformation occurs. In this study, we characterize stromal alterations that are only evident in the adult female mouse and are known risk factors for human breast cancer.

## Methods

### Animals

Animal experiments were performed in compliance with protocols approved by The Ohio State University Institutional Animal Care and Use Committee (IACUC, Protocol #2013A00000030) and in accordance with the accepted standard of humane animal care. CD1 mice (Charles Rivers, Wilmington, MA, USA) were maintained in polysulfone cages and fed a diet containing minimal levels of phytoestrogen (Harlan2019X). BPA dissolved in sesame oil was administered to pregnant mice via intraperitoneally (IP) injection as previously described (Hindman et al. 2017) unless otherwise stated in the text. Treatment with 25μg/kg • bodyweight BPA, 100µg/kg • bodyweight DES, 25µg/kg • bodyweight BPS, or equal volume sesame oil control occurred daily between embryonic days 9.5 through 18.5 (E9.5-18.5). Resulting female offspring were harvested at 4 and 12-14 weeks of age for analyses as indicated.

### Fibroblast RNA isolation and sequencing

Mammary fibroblast cells were isolated from adult mice, as previously described (Smalley 2010). Briefly, *in utero* exposed female mice were harvested between 12 and 14 weeks of age. The fourth and fifth inguinal mammary glands were excised from up to 4 litters totaling between 12-20 mice, with lymph nodes excluded, and mechanically minced to produce a fine, semi-liquid slurry. The slurry was further digested in a collagenase and trypsin mixture on a platform shaker (150 rpm at 37°C for 1 hour). The mixture is centrifuged (350x g for 3 minutes) to collect the epithelial organoids, stromal cells, fibroblasts and red blood cells in the pellet. This pellet was rinsed in red blood cell lysis buffer (Sigma-Millipore) two times and resuspended in 10% FBS DMEM. Cells were plated onto a cell culture dish for 1 hour to separate the epithelial organoids from fibroblasts. Adherent fibroblasts were maintained at 37°C/5% CO2/5% O2 until 80% confluent. Plates were washed with 1x phosphate buffered saline (PBS) and DNA and RNA harvested using the ZR-Duet DNA/RNA mini prep kit (Zymo Research). These analyses were done in three separate cohorts of mice treated with either oil or BPA *in utero*.

### Differential gene expression analysis

Sequencing reads were mapped to the MM10 genome using the STAR read aligner (Dobin et al. 2013) and transcript read counts quantified using featureCounts (Liao et al. 2014). Low expression genes with less than 10 reads across all samples were filtered out of the data set and differentially expressed genes identified using DESeq2 criteria (P≤0.05, FDR 5%)(Love et al. 2014). RNA-seq datasets are available through Geo series accession number: <u>GSE136062</u>. Statistically significant gene lists were subjected to Ingenuity Pathway Analysis (Qiagen Bioinformatics) and the ToppGene Suite (Chen et al. 2009).

### RT-PCR

Fibroblast RNA was converted to cDNA using the iScript cDNA Synthesis Kit (BioRad 170-8891) per the manufacturer’s protocol. Relative gene expression was determined as previously described using SYBR Green realtime qPCR with the primers listed in Supplemental File 1 (Patterson et al. 2015).

### Picrosirus red staining

The fourth inguinal mammary glands were harvested at either 4 weeks or 12 weeks of age, for each treatment. Mice were staged in the estrous cycle prior to harvest using vaginal cytology (Byers et al. 2012; Caligioni 2009; Cora et al. 2015), and harvested in the estrus stage. Estrus stage was confirmed through hematoxylin and eosin analysis of the dissected uterus, ovaries, and vagina from each mouse (Byers et al. 2012; Caligioni 2009). Glands were fixed in 10% neutral buffered formalin for 48 hours, transferred to 70% ethanol, and paraffin embedded. The paraffin embedded mammary glands were cut in 4um sections for picrosirius red staining. A picrosirius red staining kit (Abcam ab150681) was used to visualize collagen fibers.

Images were taken under 4x magnification with a light microscope (Nikon Eclipse 50i microscope) equipped with a camera (Axiocam color, Zeiss) and Zeiss Zen Pro software. For 4 week old mice, one image was taken per gland section, between the lymph node and nipple. For 12 week old mice, two images were taken adjacent to lymph node. Images were cropped to identical size and portions of the lymph node cut from the image. Images were randomized and blindly scored in ImageJ. Total area was then calculated of the image. A threshold was then set to select red staining within the image and the area above the threshold was calculated. Data represents the percent area above the threshold compared to total area of the image. A minimum of 6 mice from 3 separate litters was used for the dataset. A student’s t-test determined statistical significance between oil and BPA treated mice

### Hydraulic Permeability Assay

The measurement of ECM hydraulic permeability was performed with *in utero* exposed fibroblasts. Re-organization of experimentally available collagen I was performed as previously described (Hammer et al. 2017). Briefly, neutralized (NaOH) acidic rat tail type I collagen (Corning) solutions were prepared at 6mg/mL in media for the following separate experimental conditions: (1) containing oil vehicle-treated fibroblasts, (2) containing *in utero* BPA-treated fibroblasts (1800 cells/µL) and (3) no added fibroblasts for acellular conditions. Polydimethylsiloxane microfluidic columns were pre-coated with 100µg/mL fibronectin prior to collagen mixture injection, as previously determined to be optimum (Bischel et al. 2015. Polymerization of the collagen mixtures, with and without fibroblasts, was permitted prior to the addition of fresh media. Fresh media was replaced daily over 48 hours to permit the re-organization of the provided collagen by the treated fibroblasts. At this time, a height-based (1.8-2.2cm) hydrostatic pressure difference was applied between channel ports of the PDMS microfluidic device, promoting the flow of a rhodamine fluorescent fluid (used to track the flow rate) through the semi-porous collagen matrix. Hydraulic permeability was calculated using Darcy’s law (Equation 1) {Wiig, 2012 #187; Sung et al. 2009):

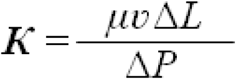

Where µ is fluid viscosity of the cell culture medium (approximated with water), v is the average fluid velocity, ΔL is the length of the PDMS microfluidic device channel and ΔP is the pressure difference across the channel due to differences in the fluid height of the two ports on either side of the device, given by (Equation 2):

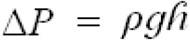

Where ρ is the density of the flow medium, g is the acceleration due to gravity (9.81m/s2) and h is the fluidic height difference between device ports.

### Mammary stiffness assay

The elastic modulus of mammary glands was determined using an unconfined compression protocol, as previously described (Mpekris et al. 2017; Papageorgis et al. 2017; Polydorou et al. 2017). Following *in utero* exposures, the fourth inguinal mammary glands were harvested from mice at either 4 weeks or 12 weeks of age, for each treatment group. Using a biopsy punch, circular sections approximately 4mm in diameter with a thickness of 1mm were excised from the fourth gland, avoiding the lymph node and excising sections towards the leading edge of the mammary gland. Sections were loaded on a mechanical testing system (Electroforce 5500, TA Instruments, Eden Prairie, MN) and compressed at a strain rate of 0.5 mm/min to a final strain of 30%. The modulus was estimated by calculating the slope of the stress-strain curve. A minimum of three sections per mouse were analyzed and averaged per data point. A minimum of 3 mice per condition from at least 3 independent litters were analyzed. A student’s t-test determined significance between oil and BPA exposed mice. A one-way ANOVA with multiple comparisons determined significance between oil, BPS, and DES.

## Results

### In utero BPA alters the transcriptome of fibroblasts in adult female mice

Several studies have implicated the mesenchymal cells surrounding the developmental mammary bud as a target of *in utero* BPA action (Hindman et al. 2017; Paulose et al. 2015; Wadia et al. 2013). In addition, the stroma plays a key role in mammary gland development and cancer risk through paracrine signaling and ECM interactions {Macias, 2012 #132}. To this end, we attempted to identify changes within the stroma that occur after *in utero* BPA exposure that may increase cancer risk. Pregnant CD1 mice were exposed to 25 µg/kg BPA or oil control E9.5-E18.5. We have previously shown this dose results in amniotic BPA levels comparable to reported human exposures (Hindman et al. 2017). Following birth, mice were aged 12-14 weeks to allow for complete epithelial ductal elongation within the mammary gland and to approximate an adult when carcinogenic risk is suspected. RNA was isolated from mammary fibroblasts and subjected to whole genome transcriptome profiling. Mice exposed to *in utero* BPA had 489 differentially regulated genes (p ≤0.05, ≥50% change) and 47 statistically significant (padj <0.05) altered transcripts when accounting for false discovery (Fig 1A, Supplemental Table 2). Interestingly, nearly all (41 of 47) of the altered genes had increased expression in the BPA exposed mice (Fig 1). Gene expression changes of these 47 genes for our 3 cohorts of mice was consistent, although very few genes showed greater than a 2-fold change in expression (Fig. 1B). Ingenuity pathway analysis of these 47 genes identified cancer, including breast, as the diseases most associated with these alterations (Table 1). In addition, genes associated with ECM and collagen regulation were the molecular functions and cellular components most enriched (Table 1).

**Table 1.** Pathways, diseases, and functions enriched amongst BPA regulated genes.

| **Molecular Function** | p-value |
| --- | --- |
| extracellular matrix structural constituent conferring tensile strength | 4.02E-08 |
| extracellular matrix structural constituent | 1.63E-07 |
| voltage-gated calcium channel activity involved in AV node cell action potential | 1.54E-05 |
| collagen binding | 3.20E-05 |
| **Cellular Component** |  |
| extracellular matrix | 4.87E-09 |
| collagen-containing extracellular matrix | 5.35E-09 |
| extracellular matrix component | 1.41E-07 |
| **Pathway** |  |
| Genes encoding extracellular matrix and extracellular matrix-associated proteins | 8.92E-10 |
| Extracellular matrix organization | 3.09E-08 |
| Integrin signalling pathway | 5.81E-08 |
| NCAM1 interactions | 1.21E-07 |
| Genes encoding core extracellular matrix including ECM glycoproteins, collagens and proteoglycans | 2.07E-07 |
| ECM-receptor interaction | 2.54E-07 |
| Genes encoding collagen proteins | 2.97E-07 |
| Collagen chain trimerization | 4.17E-07 |
| Collagen biosynthesis and modifying enzymes | 3.11E-06 |
| **Disease** |  |
| Colorectal cancer | 9.22E-10 |
| Breast or pancreatic cancer | 9.99E-10 |
| Breast or ovarian cancer | 1.39E-09 |
| Development of vasculature | 1.73E-09 |
| Angiogenesis | 1.89E-09 |
| Breast or gynecological cancer | 6.34E-09 |
| Female genital tract cancer | 8.78E-09 |
| Dupuytren contracture | 1.03E-08 |
| Breast or ovarian carcinoma | 1.38E-08 |

**Figure 1.**
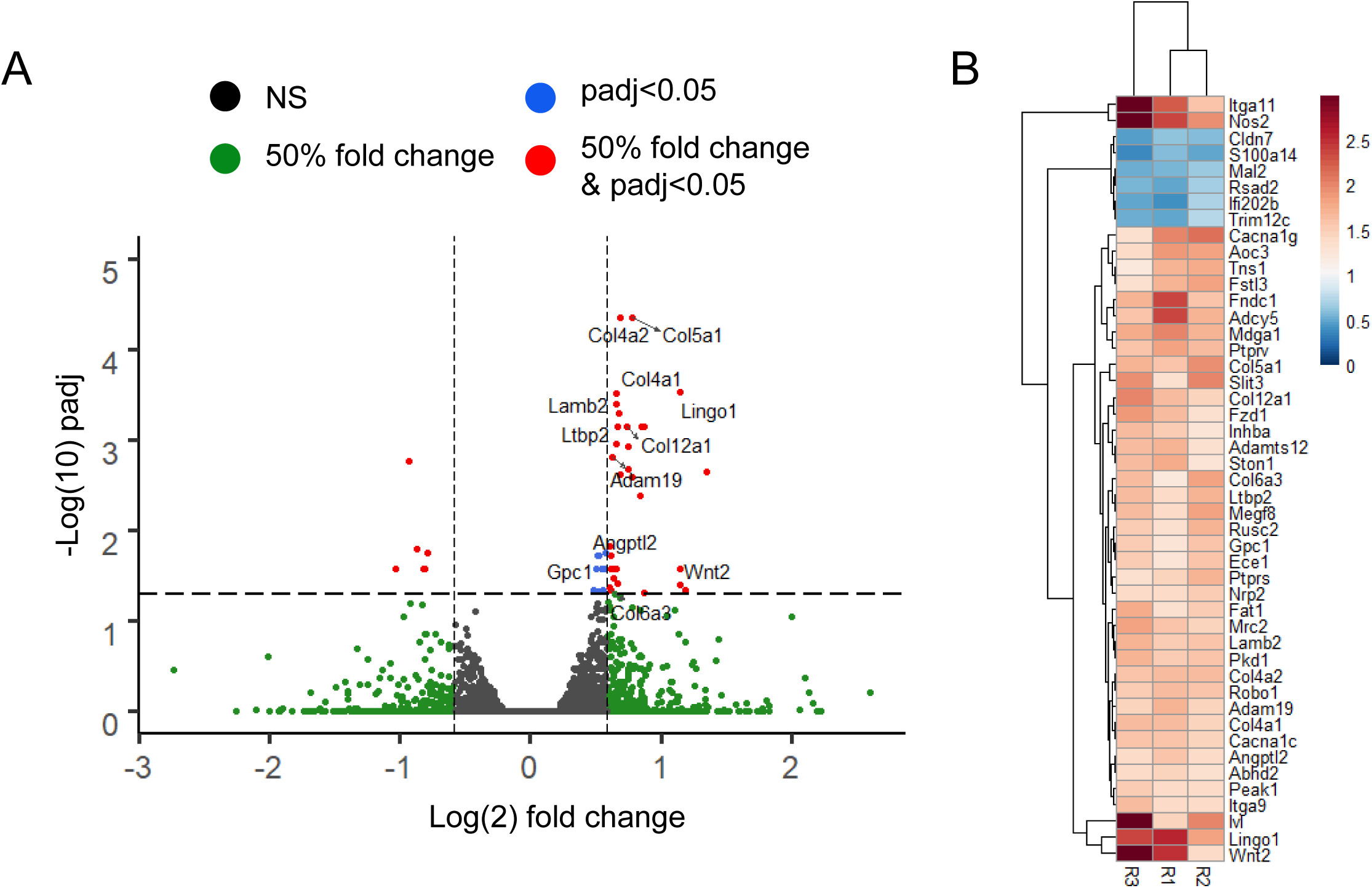
*In utero* BPA exposure drives long term changes in mammary fibroblast gene expression. Fibroblasts were isolated from the mammary glands of 12-week old female mice exposed *in utero* to 25 μg/kg • bodyweight BPA or oil control from E9.5-18.5. RNA from the fibroblasts was sequenced and analyzed for differential gene expression. **A**. A Volcano plot is depicted of all analyzed genes with cutoffs for fold change, p-value, and p-adjusted (padj) value depicted. **B**. A heatmap of the 47 genes with a p-adjusted value ≤ 0.05 is depicted illustrating the fold change in each of the three cohorts of mice used for the transcriptome analysis.

Changes in collagen deposition have been proposed as a risk factor for breast cancer, so we investigated the collagen family of genes in more detail. A total of 15 collagen genes demonstrated a change in expression (p≤0.05) in BPA exposed mice. In fact, increased collagen expression in BPA exposed mice were observed in our 3 cohorts (R1, R2, R3) especially amongst the most highly expressed collagen genes (Fig. 2A). We thus validated these changes amongst an additional 3 cohorts of 12-14 week old mice whose mothers were exposed to BPA or oil via oral gavage, as opposed to IP injection. In these additional samples, we again see a general increase of 20-50% expression across the most highly expressed collagen genes when exposed *in utero* to BPA (Fig. 2B). In line with previous studies, the route of exposure does not effect this phenotype (Taylor et al. 2008).

**Figure 2.**
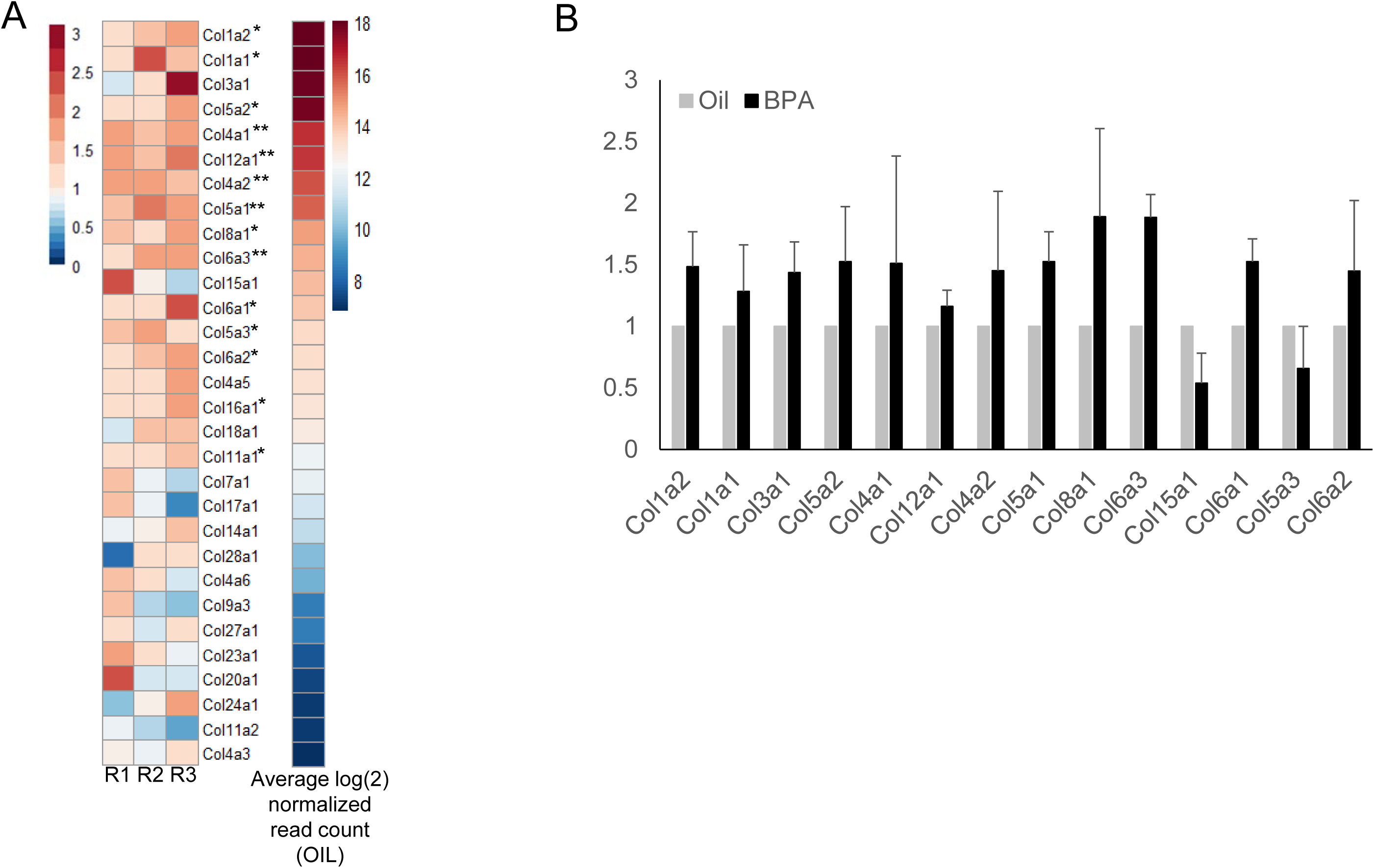
Collagen genes are more highly expressed in BPA exposed mammary glands. **A**. A heatmap showing the RNA expression of the most abundatntly epxressed collagen genes is depicted. RNA was isolated from fibroblasts of 12 week old female mice in three separate cohorts. Collagen genes are ordered from top to bottom based upon the average normalized read counts in the control samples. **B**. Collagen genes that had increased expression in our RNA-seq data set were validated on an additional 3 mice cohorts using q-PCR. Average normalized expression with standard error is depicted.

### BPA exposure increases collagen deposition in adult female mice

In order to determine if these transcriptional changes are resulting in increased collagen deposition in adult mice, we stained mammary glands of exposed and control mice with picrosirius red. We performed these analyses early in mammary development at 4 weeks of age and again after complete ductal elongation at 12 weeks of age. At 4 weeks of age, we see no change in collagen expression between BPA and control mice (Fig 3A). However, at 12 weeks of age, BPA exposed mice have significantly more area of the mammary gland stained for collagen deposition (Fig 3B).

**Figure 3.**
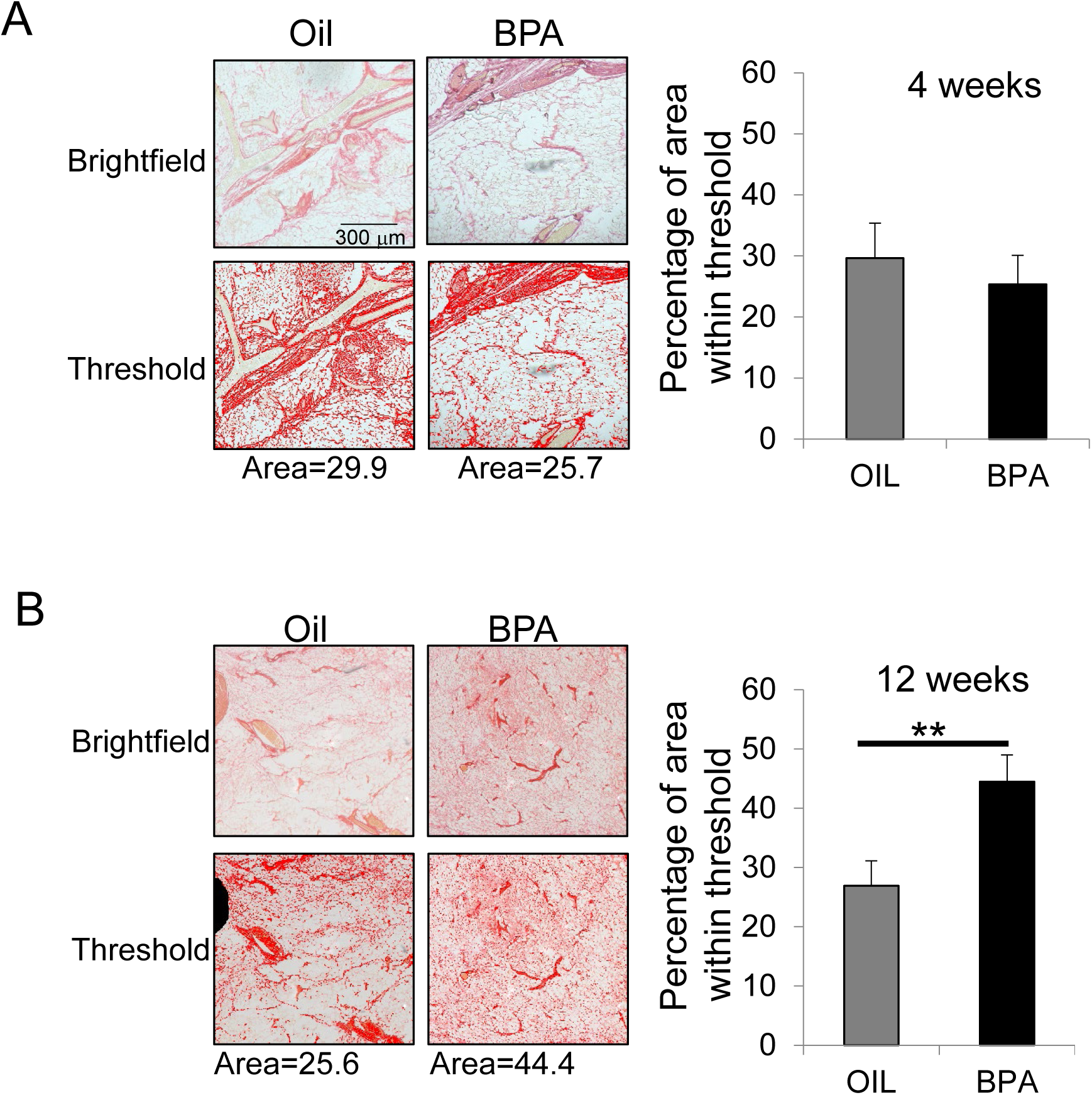
BPA exposed mammary glands have increased collagen deposition in adult female mice. **A**. Mammary glands from 4-week old mice exposed to BPA or oil *in utero* were stained with picrosirius red to visualize collagen. Total collagen staining was quantified by the area of red staining in images (right graph). A representative brightfield image and thresholded image used for quantification are depicted. Selected images represent the data point closest to the median for that sample set. **B**. Mammary glands from 12-week old mice quantified as in (**A**) for total area of the mammary gland stained for collagen. ** represents p≤0.01.

### In utero BPA increases collagen matrix density and gland stiffness

Collagen expression and breast density are significant risk factors for breast cancer. Thus, we aimed to determine if stromal reprogramming is increasing these known risk factors. We first tested the ability of fibroblasts to remodel the ECM *in vitro*. Isolated fibroblasts from 12-14 week old mice exposed to either BPA or oil control were plated with collagen in a microfluidic column. After allowing 48 hours for fibroblasts to remodel the collagen matrix, a fluorescent liquid was pumped through the column. Using the flow rate and pressure required to pump the liquid, a permeability constant (κ) indicative of the collagen matrix density can be calculated (Fig 4A). BPA exposed mice had decreased collagen permeability as compared to the oil exposed fibroblasts (Fig 4B). This decreased permeability indicates that BPA exposed fibroblasts created a denser collagen matrix than oil exposed fibroblasts and no fibroblasts (control).

**Figure 4.**
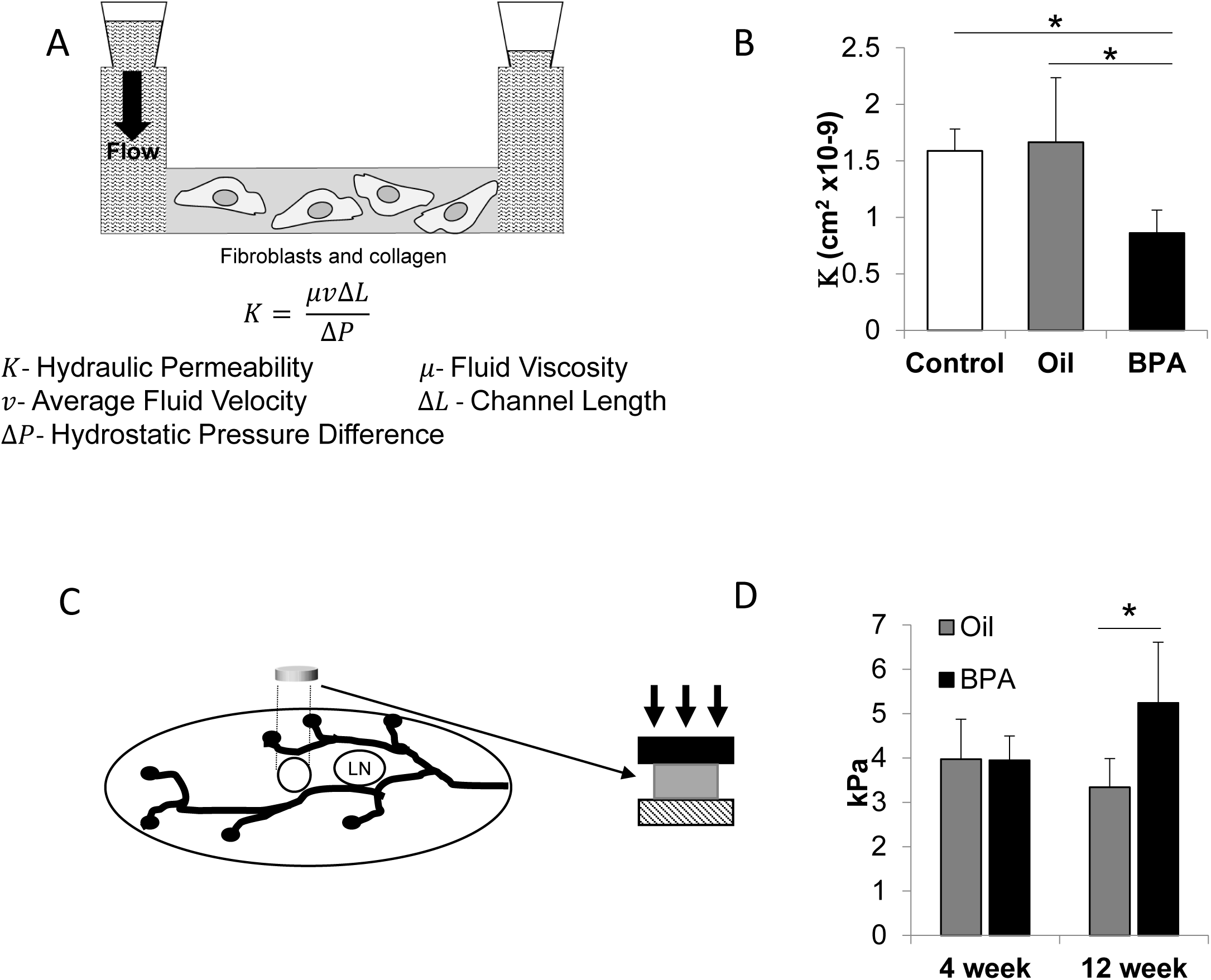
BPA decreases collagen matrix permeability and increases mammary gland stiffness. **A**. An *in vitro* assay was utilized to measure fibroblast organization of a collagen matrix. A PDMS microfluidic column was pre-coated with fibronectin and then fibroblasts isolated from 12 week old mice were plated with collagen. After 48 hours of fibroblast-mediated remodeling of the collagen matrix, a fluorescent liquid was pumped through the column. Measurement of the pressure and flow rate was used to calculate the permeability (κ) of the collagen matrix arranged by the fibroblasts. **B**. Mammary fibroblasts were isolated from BPA and control 12-week old mice and plated within the column as depicted in (**A**). BPA exposed fibroblasts generate an ECM with decreased permeability compared to oil control. Control represents the permeability of collagen with no cultured fibroblasts. **C**. The diagram depicts the methodology to measure mammary gland stiffness. A circular section approximately 4mm diameter and 1 mm thick is removed adjacent to the lymph node from mammary glands. The amount of pressure required to compress these discs is measured to determine the stiffness of the gland. **D**. Mammary glands from 4- and 12- week old mice were analyzed as described in (**C**) for the stiffness. 4-week old mammary glands show no difference in stiffness between BPA and oil treated mice. At 12 weeks of age, mice exposed *in utero* to BPA show a significant increase in gland stiffness compared to oil controls. * represents p≤ 0.05.

We next aimed to determine if these fibroblast-mediated changes affect mammary density and stiffness *in vivo*. We thus adapted a tumor stiffness approach to measure mammary gland stiffness (Fig 4C). Mice treated *in utero* with BPA or oil by oral gavage were analyzed for changes in gland stiffness. In line with our transcriptome and *in vitro* analyses, adult 12 week old mice have increased mammary gland stiffness as opposed to control mice. Interestingly, 4 week old mice did not demonstrate any change in mammary gland stiffness between BPA and control mice (Fig 4D).

As BPA exposure increases ECM density and gland stiffness, which are associated with increased breast cancer risk, we wanted to determine if this phenotype is conserved in response to other estrogenic EDCs. To this end, we generated an additional cohort of mice that were exposed *in utero* to DES, a strong estrogen known to increase breast cancer risk in humans, and BPS, a very weak estrogen used in BPA-free products. Pregnant mice were exposed from E9.5-E18.5 with 100µg/kg • bodyweight DES, 25µg/kg bodyweight BPS, or oil control via oral gavage. Doses were chosen to be consistent with previous literature in mouse studies or known human exposures (Hilakivi-Clarke et al.2013; Kolla et al. 2018; LaPlante et al. 2017; Tucker et al. 2018). At 12 weeks of age, mice were subjected to our breast stiffness assay. DES exposed mice demonstrated a significantly increased mammary gland stiffness while the weak estrogen, BPS, was not statistically different from our oil control (Fig 5). These data suggest that estrogenicity may be the driving factor of this phenotype and demonstrate that the one EDC known to increase breast cancer risk in humans (DES) drives stromal changes associated with the disease.

**Figure 5.**
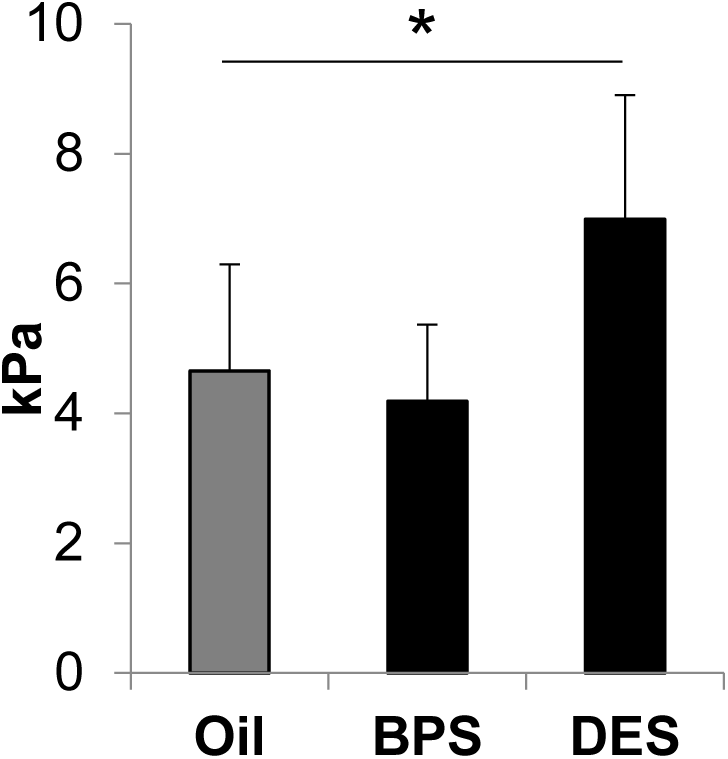
DES but not BPS exposures recapitulate the increase in breast stiffness of BPA exposures. A cohort of mice were treated *in utero* with either 100µg/kg • bodyweight DES, 25µg/kg • bodyweight BPS, or oil control between E9.5-18.5. At 12 weeks of age, mammary glands were harvested and gland stiffness measured as depicted in Figure **4C**. BPS showed no change, but DES showed a significant increase in gland stiffness when compared to oil control. * represents p≤ 0.05.

## Discussion

### BPA induces adult phenotypes associated with human breast cancer risk factors

In this study, we focused our efforts to analyze the changes within the stroma induced by *in utero* BPA exposures that persist into adulthood when carcinogenic transformation occurs. We identified fibroblast-specific reprogramming of the transcriptome associated with ECM organization. These changes are highlighted by increased collagen deposition, resulting in a denser and stiffer mammary gland. Breast density is one of the strongest risk factors for breast cancer and numerous models have demonstrated that altering the extracellular matrix alone can contribute to oncogenesis (Schedin and Keely 2011). In fact, mutation of the MMP cleavage site in Col1a1 gene results in increased collagen density and susceptibility to mammary carcinogenesis (Provenzano et al. 2008). Further, animal and *in vitro* studies have demonstrated that increasing ECM density drives a malignant phenotype (Chaudhuri et al. 2014; Schedin and Keely 2011). These data are supported by a human study which correlated breast stiffness to cancer risk (Boyd et al. 2014). Taken together, our data provides a specific EDC phenotype that has been directly linked to breast cancer risk in humans.

Early life EDC exposure has life-long consequences towards human health. Interestingly, we found that these phenotypes of increased ECM density and gland stiffness, known to be risk factor for breast cancer, were not evident in young mice. These data support the idea that carcinogenic risk progresses throughout life and the impact of these exposures is not necessarily apparent at the time of exposure. In line with this hypothesis is the fact that DES daughters do not display increased breast cancer risk until the women have reached 40 years of age ((NIH) 1999; Palmer et al. 2006).

### BPA alternatives and the importance of estrogenic activity

Identification of new EDCs is mostly driven by the ability to drive a hormone response in various *in vitro* and *in vivo* assays. While the ability of an EDC to activate the ER is associated with potential risk, it is unclear if it is the estrogenic action of these compounds that drives cancer risk. Furthermore, much of the analysis of BPA and its alternatives have used surrogate phenotypes other than tumor formation to implicate cancer risk. These phenotypes include developmental abnormalities such as increased ductal branching and changes in the terminal end bud number (Hindman et al. 2017; Markey et al. 2001; Moral et al. 2008; Munoz-de-Toro et al. 2005). However, these phenotypes are not universally observed in all strains of mice. For example, the FVB strain shows none of these developmental phenotypes following *in utero* BPA exposure, yet this strain still displays increased cancer susceptibility when challenged with a chemical carcinogen (Weber Lozada and Keri 2011). Increased ductal branching and epithelial ducts would contribute to an increased mammographic breast density which is associated with breast cancer. However, mammographic breast density, a measure of the percent glandular tissue in the gland, is not higher in DES daughters (Strohsnitter et al. 2018). That finding suggests that certain developmental phenotypes being assessed in mice don’t carry over to human biology. Together, these data suggest that many of the surrogate phenotypes are not great predictors of risk. It is further unclear if specific estrogenic activity or affinity is also predictive.

In our system, we show that BPA and DES demonstrate increased gland stiffness in the adult animal and this is a phenotype that is associated with cancer risk in humans. However, BPS did not show any change in stiffness. Interestingly, BPS induces many of these developmental abnormalities in the mammary gland which are seen with BPA and are typically used as a measure of cancer risk (Kolla et al. 2018). While no study has directly tested BPS in a cancer susceptibility model, such as DMBA, its association with these phenotypes has raised concerns. Our data suggests that BPS may indeed be associated with less breast cancer risk when compared to BPA. It is worth noting that BPS has weaker estrogenic activity than BPA, suggesting that activity may be the driver of risk (Kojima et al. 2019; Ng et al. 2015; Rochester and Bolden 2015). In contrast, DES is a very strong estrogen and similarly drove a strong gland stiffness phenotype. Further analysis on a wider range of estrogenic compounds at a range of dosages will be needed to determine if estrogenic activity is in fact the primary driver of this phenotype and cancer risk.

